# Experience-dependent sex differences in the role of dorsal striatal dopamine D1 receptor activity in methamphetamine self-administration revealed by a novel TREND model

**DOI:** 10.1101/2024.07.06.601793

**Authors:** InduMithra Madhuranthakam, Martin O Job

## Abstract

**BACKGROUND:** The role of dorsal striatal dopamine D1 receptor systems in the mechanism of methamphetamine self-administration (METH SA), and sex differences in this role, are unclear. We hypothesized that this role would be sex and METH experience-dependent. Because prior experience regulates subsequent effects of drugs, we developed a novel model to account for this interaction, termed the TREND model (Time-Related-Experience-Normalized-Dynamics) for drug SA analysis. We tested our hypothesis by comparing results from the new TREND model and the current model.

**METHODS:** For model validation, we reanalyzed previous data (Job et al., 2020) with the aim of determining which model (current or TREND) was more effective as an analytical tool. We compared variables from each model with the effect of Clozapine-N-Oxide (CNO, chemogenetic ligand) on METH SA. We employed regression analysis, median split, ANOVA to see which could reveal sex and experience dependency of dorsal striatal dopamine D1 receptor system.

**RESULTS:** The current model variables were unrelated to CNO effect, with no sex differences in these relationships. TREND model revealed new variables that were unrelated to current variables but related to CNO effect on METH in males and females, with sex differences in these relationships. TREND, but not the current model, detected sex differences when comparing males and females with prior high, but not low, behavioral response variables.

**CONCLUSIONS:** TREND model is more sensitive than the current model for detecting experience-dependent sex differences in the role of the dorsal striatal dopamine D1 receptor systems in the mechanism of METH SA.

## Introduction

There is an epidemic of methamphetamine (METH) use disorders (MUD) with huge socioeconomic impact and an urgent need for pharmacotherapeutic intervention strategies. There are sex differences in METH use and abuse, and the consequences thereof (Daiwile et al., 2022). To address this health crises, we need to understand the mechanism governing METH consumption and sex differences in METH consumption. To achieve these goals, more sensitive tools are required, including novel models for obtaining important data that may be underestimated by current models.

Given the evidence that 1) dopaminergic mechanisms in the dorsal striatum are thought to play an important role in the progression of drug experience from recreational to habitual drug use (Belin and Everitt, 2008; Dalley and Everitt, 2009; Everitt and Robbins, 2005; Lüscher et al., 2020; Vanderschuren and Everitt, 2004; Veeneman et al., 2012), it is not farfetched to suggest that inhibition of dorsal striatum dopamine D1 receptor might alter METH self-administration (SA) in a METH experience-dependent manner (Fig 1). The dorsal striatum is highly regulated by dopamine D1 receptor systems. For example, it exhibits high levels of D1 receptor expression (Dubois et al., 1986; Savasta et al., 1986; Shishido et al., 1997; Somkuwar et al., 2016), and METH SA increases dopamine D1 receptor expression in the dorsal striatum (Shishido et al., 1997; Somkuwar et al., 2016), though not all studies agree (Okita et al., 2018; Tong et al., 2003). Additionally, METH SA has been shown to interact specifically with several neurobiological systems that interact with dopamine D1 receptor (Wang and McGinty, 1995; Yoshida et al., 1995). Further, there are suggestions that the dorsal striatum is significantly altered by prolonged psychostimulant use (Dalley and Everitt, 2009; Everitt and Robbins, 2013, 2005; Lüscher et al., 2020; Tong et al., 2003).

**Figure 1.**
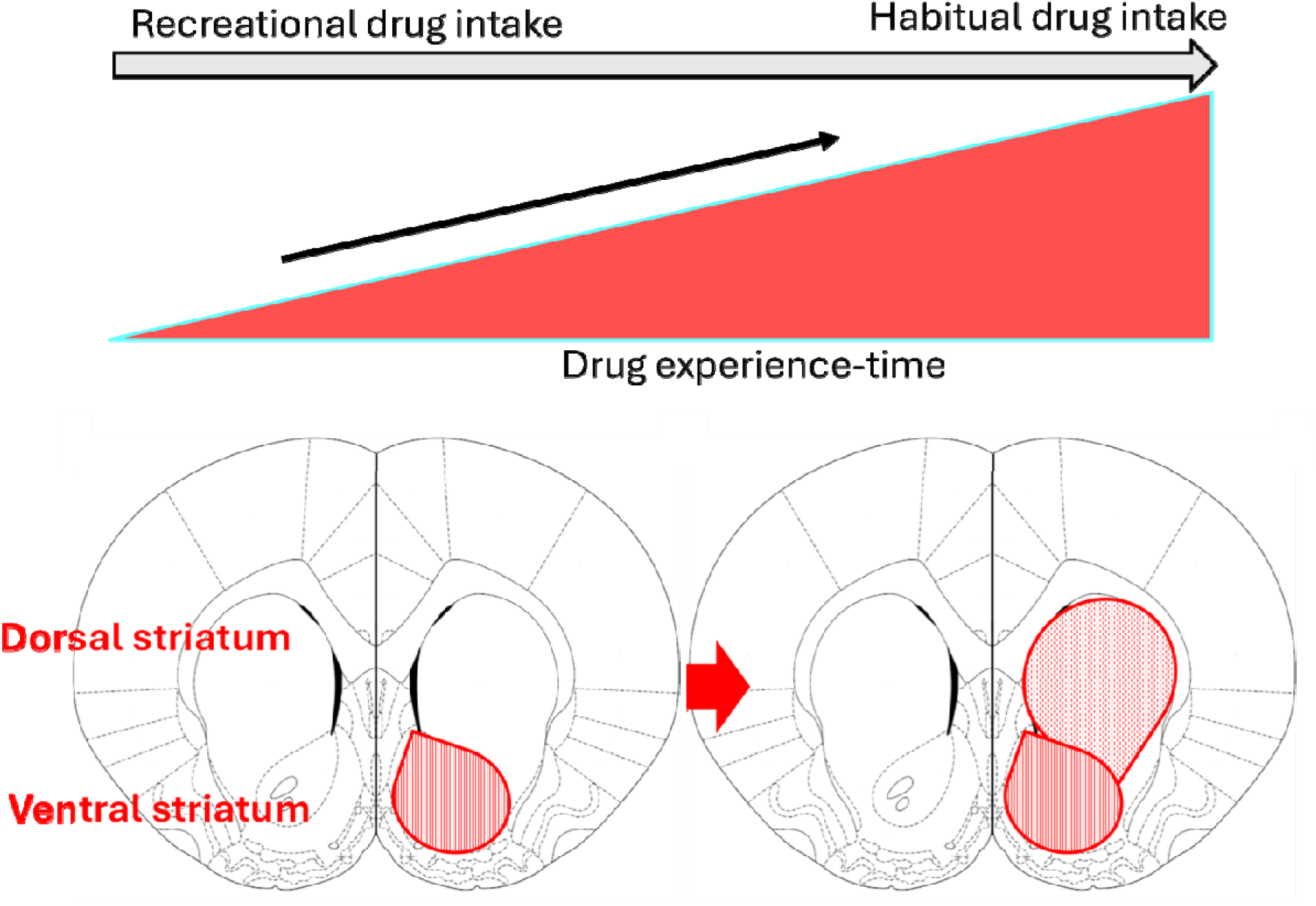
Experience-dependent engagement of the dorsal striatum. As subjects progress from recreational drug use to habitual drug use, there is thought to be an increase in the engagement of the dorsal striatum in the mechanism of drug use behavior. This can be conceptualized as an experience-dependent recruitment of the dorsal striatum. The implication is that the role of dopamine D1 receptor systems in the dorsal striatum will be experience-dependent.

However, studies examining the impact of blocking dopamine D1 receptors in the dorsal striatum on METH SA have yielded inconsistent results. Specifically, some research suggests a decrease in METH SA following D1 receptor blockade (Avchalumov et al., 2020), while other studies indicate an increase (Kreisler et al., 2020; Oliver et al., 2019) or report no observable changes (Job et al., 2020). A careful study of these reports suggests that we cannot rule out differential METH experience prior to the tests to determine the effects of dorsal striatum dopamine D1 receptor on METH SA as the reason for the paradoxical effects observed (Table 1), but this is not clear. In addition to the question of METH experience-dependency, there is also the question of sex differences. We previously reported no sex differences in the role of dorsal striatal D1 receptor systems in METH SA (Job et al., 2020).

It is possible that the paradoxical results in Table 1 and inconsistencies in the observation of sex differences in METH SA (Bachtell et al., 2023; Daiwile et al., 2022, 2021; Holtz et al., 2012; Job et al., 2020; Johansen and McFadden, 2017; Kearns et al., 2022; Lewandowski et al., 2023; Lin et al., 2024; Pena-Bravo et al., 2019; Pittenger et al., 2021; Venniro et al., 2017; Westbrook et al., 2020) are due to drug self-administration model sensitivity issues. By this we mean that the drug self-administration models currently used in the field, though widely used, may not be as effective as we believe in obtaining behavioral variables that are sensitive enough to reveal experience-related and/or sex-related differences in METH SA. Resolving this limitation is crucial for understanding the mechanisms underlying METH SA.

To resolve this, we need to account for the relationship between prior drug experience and subsequent drug consumption. Prior experience with METH influences the responsivity of subsequent METH SA to pharmacological regulation. For example, there is evidence that pre-exposure to METH has a modulatory effect on later METH SA, including the dopaminergic mechanisms that govern such behavior (Chojnacki et al., 2020; Culbertson et al., 2009; Kazahaya et al., 1989; Lorrain et al., 2000; Mcfadden et al., 2013; McFadden et al., 2015, 2012; Segal et al., 2003; Vezina et al., 1999). Further, the level *per se* of prior METH experience can differentially modulate subsequent METH SA response: a comparison of METH experienced rats trained under restricted access versus extended access revealed that subjects taking METH under extended access conditions were more sensitive to the effects of aripiprazole, a dopamine D2 receptor partial agonist, on METH SA (Wee et al., 2007). Additionally, varying levels of prior METH exposure have been shown to result in differential sensitivity to the inhibitory effects of dopamine D3 receptor partial agonists on METH SA (Orio et al., 2010).

## Table

| References | (Oliver et al., 2019) | (Job et al., 2020) |
| --- | --- | --- |
| Effect of DSD1R inhibition on METH SA | Increase | No effect |
| Dose of CNO (mg/kg i.p.) | 5.0 | 1.0 |
| Dose of METH SA (mg/kg/infusion) | 0.05 | 0.1 |
| Number of days of METH SA during training | 10 – 11 days | 20 days |
| Duration of METH SA/day (h/day) | 6 | 6 |

*Differences in prior METH experience may be related to the differential effects of dorsal striatal D1 receptor activity inhibition on METH intake.*
*Both studies cited above (Job et al., 2020; Oliver et al., 2019) used the same animal model (rats), chemogenetic ligand (CNO, Clozapine-N-Oxide) and duration of METH SA per day (6 h). However, these studies used different doses of METH for the SA protocol and different number of days of METH SA before the assessment of CNO-mediated inhibition of DSD1R expressing neurons on METH SA: the rats in the study by (Oliver et al., 2019) received METH at a dose of 0.05 mg/kg/infusion for 10-11 days whereas the rats in the study by (Job et al., 2020) received a dose of 0.1 mg/kg/infusion for 20 days. Chemogenetic-mediated inhibition of dorsal striatal dopamine D1 receptor led to increases (Oliver et al., 2019) and no effect (Job et al., 2020) in METH SA. A thorough analysis suggests that the rats in the study by (Oliver et al., 2019) have less METH experience (dose x number of sessions) than those in the study by (Job et al., 2020). Thus, the increases (Oliver et al., 2019) and no effect (Job et al., 2020) in METH intake following dorsal striatal dopamine D1 receptor inhibition may be related to differences in METH experience prior to the test.*

Moreover, evidence suggests that the efficacy of systemic D1 receptor blockade in regulating amphetamine SA is contingent upon an individual’s history with amphetamine. In a study, the D1 receptor antagonist, SCH23390 exerted differential effects on amphetamine SA in subjects that had no prior experience with amphetamine versus subjects with prior experience with amphetamine (Pierre and Vezina, 1998). Notably, the inhibitory effectiveness of systemic SCH23390 was found to correlate with the extent of previous amphetamine exposure, being ineffective in subjects with low prior experience and inhibitory in those with high experience (Pierre and Vezina, 1998).

From the above cited references, including careful analysis of Table 1, we hypothesize that METH SA in subjects with low and high prior METH experience will be differentially responsive to the inhibition of the activity of dorsal striatum D1 receptor-expressing neurons.

To be able to test this hypothesis, we need a better model than the current model being utilized in the field. This new model should account for the interaction between prior experience with a drug and subsequent drug taking behavior in a dynamic way. A mathematical expression of such a variable(s) would be as below:

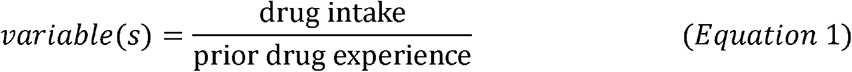

When we consider that prior drug experience includes all the drug that the subject has taken up until the point of observation of the drug intake in equation 1 above, but will also include the drug intake at the moment of observation, this equation can be rewritten as follows:

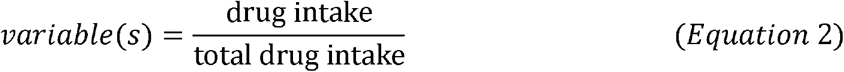

Because drug intake is not taking place over time or sessions, total drug intake can be expressed as cumulative drug intake. We therefore conceptualize that drug intake by the subject is occurring on top of cumulative drug intake over time. The variable(s) that we are trying to assess would therefore be drug intake as a function of cumulative drug intake:

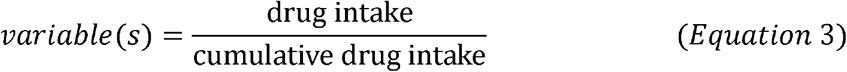

The requisite variable(s) can be obtained from the combination of graphs of a daily drug intake over time (Fig 2A) and cumulative drug intake over time (Fig 2B). The time course of drug intake/ cumulative drug intake, when obtained, can be fit with an exponential curve function (Fig 2C). The exponential function is as below:

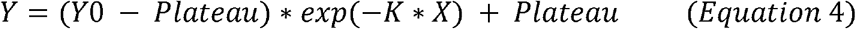

**Figure 2.**
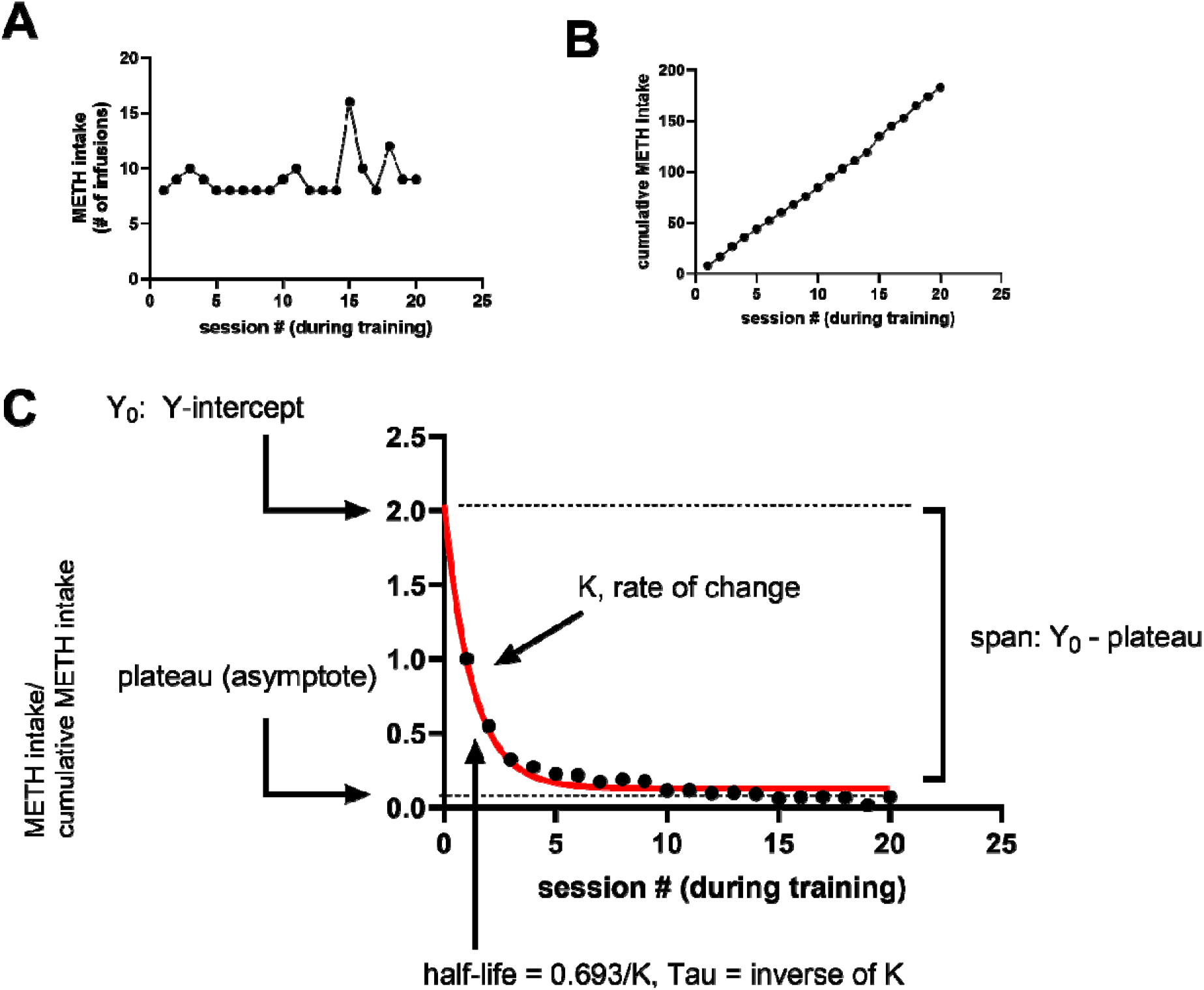
Introducing the new TREND model. Fig A shows a graph of drug self-administration design with 1 session per day for 20 sessions. The y-axis is the number of responses, or response rate, or number of reinforcers earned. Fig B is like Fig A but is a cumulative plot of the responses in A. Fig C is the TREND model and while we plot session # on the x-axis, the y-axis is a plot of drug intake/cumulative drug intake – note that on the first day the value of y = 1. This is because on the first day the drug intake is the same as the cumulative drug intake. TREND is fitted with an exponential decay function, see equation 4-5. The variables derived from TREND include Y_0_ (the y-intercept, the drug intake/cumulative drug intake at session # = 0, plateau is the asymptote (see Fig C), span is the difference between Y_0_ and the plateau. K is the rate of change. See other variables (half-life, Tau).

Where the y-intercept (Y_0_), decay rate (K), half-life and tau (both functions of K), plateau (or asymptote) and span (which is derived from Y_0_-plateau, see variables in Fig 2C). The equation can also be expressed as:

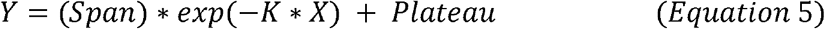

Equations 1-5 represent the TREND model which means Time-Related Experience-Normalized Dynamics. Drug intake is experience-normalized because we conceptualize that it is occurring over time on a continuum rather than as an isolated behavior.

We developed a new model because we felt that the current model is missing the interaction between drug intake and what we term drug experience-time complex (cumulative drug intake progressing on a time continuum) (Figure 2). This new model integrates drug intake and the drug experience-time complex to give rise to an exponential model with new variables (Figure 2). The current model yields variables such as escalation, total drug intake over time while the TREND model yields the variable(s) mentioned above.

We decided to test our hypothesis using both the current model and the new TREND model. To compare these models (current models versus TREND model), we re-analyzed previous data we published (Job et al., 2020) where we reported no effect of chemogenetic-mediated inhibition of neurons in the dorsal striatum that express dopamine D1 receptors on METH intake in a drug SA paradigm, with no sex differences observed for male and female Long Evans rats.

## Methods and Materials

This is a reanalysis of data as presented in a previous publication (Job et al., 2020) but using an entirely novel approach (this is the first time this is being employed in the field). The previous experiment included data for the effects of CNO on saline SA and METH SA. We reanalyzed data for only the subjects that self-administered METH. For this current analysis, a total of twenty-seven (n = 27) subjects were employed (n = 9 males and n = 18 females).

Animals: All procedures and treatments were approved by the National Institute on Drug Abuse Animal Care and Use Committee and followed the guidelines outlined in the National Institutes of Health (NIH) *Guide for the Care and Use of Laboratory Animals*. The animals, female transgenic Long Evans (Drd1a-iCre#3), were obtained from the National Institute on Drug Abuse (NIDA) Transgenic Rat Project. Rats were single housed on a 12-hour reversed light/dark cycle with free access to food and water. Stereotaxic surgery for DREADD-adeno-associated virus (AAV) injections and jugular vein catheterizations: For details regarding stereotaxic surgical techniques for the type of DREADDs (Designer Receptors Exclusively Activated by Designer Drugs) including the adeno-associated virus type (AAV1-hSyn-DIO-hM4D (Gi)-mCherry serotype 1) and the delivery of DREADDs into the dorsal striatum, see (Job et al., 2020). Briefly, stereotaxic surgical techniques, under aseptic conditions, were employed for chemogenetic manipulations using <u>D</u>esigner <u>R</u>eceptors <u>A</u>ctivated by <u>D</u>esigner <u>D</u>rugs (or DREADDs) technology.

The experimental design is described in Job et al (2020). The DREADDs-containing adeno-associated virus (AAV) (AAV1-hSyn-DIO-hM4D (Gi)-mCherry serotype 1) were obtained from NIDA Genetic Engineering and Viral Vector Core (GEVVC). Rats were injected via the intracranial route to deliver 1.0 μL of a solution containing viral titer 9.57 × 10^11^ VG/mL to the dorsal striatum at the following coordinates (from Bregma): A/P +1.6, M/L ±3.0, D/V −5.0. The AAV injection was done over 5 min using 10-μL Hamilton syringes (Hamilton, Reno, NV) driven by a microsyringe pump attached to an automated controller (World Precision Instruments, Sarasota, FL). On the same day as the intracranial injection, catheters were placed in the jugular vein. For jugular vein catheterization surgery, see (Job et al., 2020). All animals were allowed to recover from surgery for approximately one week before METH SA procedures were initiated.

Self-administration and chemogenetic manipulations: After rats had recovered from the surgical procedures described above, the rats were allowed to self-administer METH (0.1 mg/kg/infusion, or saline in control rats) for 6 h per session for 20 days as described in Job et al (2020). We defined METH or saline SA as the number of active lever presses for drug. Thereafter the pretreatment experiments (counterbalanced design) were conducted. At this time, the pretreatments, vehicle (VEH) or CNO (1 mg/kg), were injected via the intraperitoneal route 30 min before METH SA sessions. The SA experiments were done every day of the week, but the tests (pretreatment of CNO and/or VEH) were conducted for four days of the week (the day immediately after the weekend recess was not included because the SA on this day can be very variable). We conducted tests in 4 weeks with the experiments conducted in counterbalanced design: rats that had received CNO previously now received VEH injection, and vice versa, with a week of no injection in between these treatment weeks. After the experiments, the rats were euthanized and the injection sites of the AAVs were verified to be in the dorsal striatum. For more experimental details, see (Job et al., 2020).

### Variables

The variables for the current model included escalation. Escalation was calculated using linear regression analysis of the METH intake-time course, with this variable being the slope (see Fig 2A). Another variable was cumulative drug intake during the 20 days of training (Fig 2B). The variables for the TREND model include Y_0_, plateau, K, half-life, Tau and span (Fig 2C, see equation 4-5).

Another variable estimated was CNO effect on drug intake: We calculated the CNO effect (representing dorsal striatal dopamine D1 receptor activity inhibition on total METH intake level per appropriate test week) relative to VEH effect on METH SA as follows:

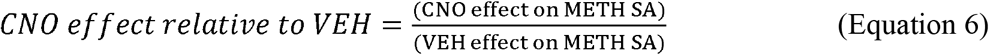

We log transformed this variable for analysis.

The goal of this study is to determine if variables from the current and TREND model can predict CNO effect, and to characterize which of these models was more sensitive to detect sex differences.

Statistical analysis: GraphPad Prism v 9 (GraphPad Software, San Diego, CA), SigmaPlot 14.5 (Systat Software Inc., San Jose, CA) and JMP Pro v 16 (SAS Institute Inc., Cary, NC) were employed for statistical analysis. Statistical significance was set at P < 0.05 for all analyses with appropriate post hoc test employed when significance was detected.

Model comparison: We compared males and females for average variables using unpaired t-tests (Fig 3). We employed regression analysis to determine the relationships between variables and compared the relationship (slope) between males and females (Fig 3). Because the level of prior METH experience may result in an experience-dependent role of dorsal striatum dopamine D1 receptor systems, we conducted experiments to compare the current model with the TREND model to determine which would yield variables that reflect this experience-dependency. For the current model, we chose cumulative METH intake. We conducted a median split analysis on male and female subjects for the cumulative METH intake to designate a high METH experience versus low experience group. For the TREND model, we conducted median split analysis on the span to designate high span and low span males and females. These procedures were followed by Two-way ANOVA with factors as SEX (males and females) and group (high versus low responders) to determine if we would detect a SEX × group interaction.

**Figure 3.**
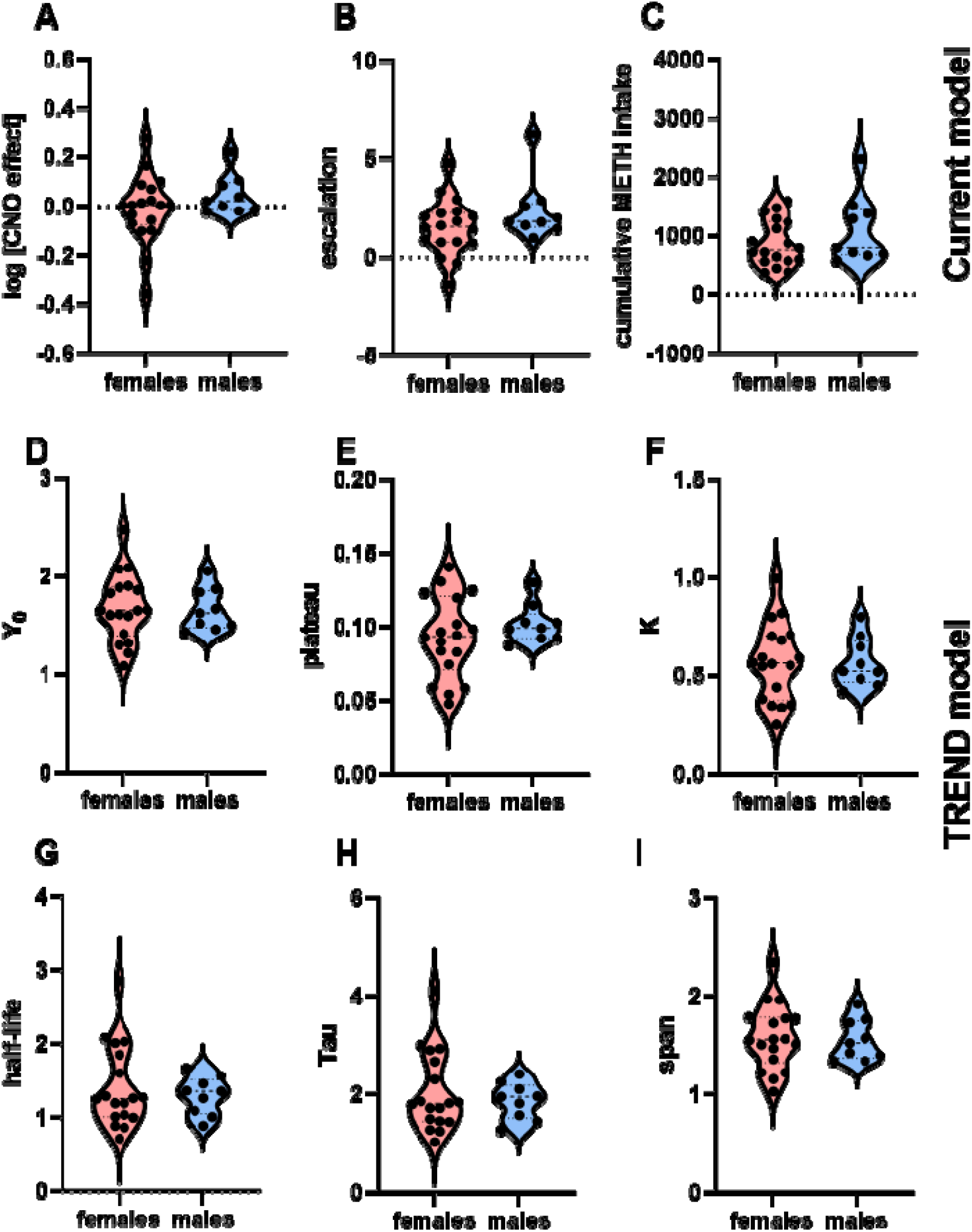
No sex differences detected for comparisons of variable averages. We did not detect any sex differences (unpaired t-tests) for the effect of CNO (Fig A), and in the variables for the current model such as escalation (Fig B) and cumulative METH intake (Fig C). We did not detect any differences between sexes for all the variables derived from the TREND model (Fig D-I).

## Results

There were no detected sex differences in average CNO effect (P = 0.2724, Fig 3A), as reported previously (Job et al., 2020). Unpaired t-tests were conducted to determine if there were significant differences between males and females for escalation and cumulative METH intake (current model variables). There were no sex differences for escalation (P = 0.1709, Fig 3B) and cumulative METH intake (P = 0.2238, Fig 3C). For TREND model, unpaired t-tests were conducted to determine if there were significant differences between males and females for Y_0_, plateau, K, half-life, Tau and span. There were no sex differences for Y_0_ (P = 0.8960, Fig 3D) plateau (P = 0.3444, Fig 3E), K (P = 0.9938, Fig 3F), half-life (P = 0.5591, Fig 3G), Tau (P = 0.5588, Fig 3H), and span (P = 0.8295, Fig 3I).

For individual females, the current model variables escalation (P = 0.4156, Fig 4A) and cumulative METH intake (P = 0.8026, Fig 4B) were not correlated with CNO effect. The variables escalation (P = 0.6578, Fig 4C) and cumulative METH intake (P = 0.2845, Fig 4D) were not correlated with CNO effect for individual males also. However, for females, the TREND model variables (Y_0_, K and span) could predict CNO effects (P < 0.05, Fig 4E, G, J). Similarly for males, Y_0_, and span could predict CNO effects (P < 0.05, Fig 4K, P) with K approaching significance (P = 0.055, Fig 4M).

**Figure 4.**
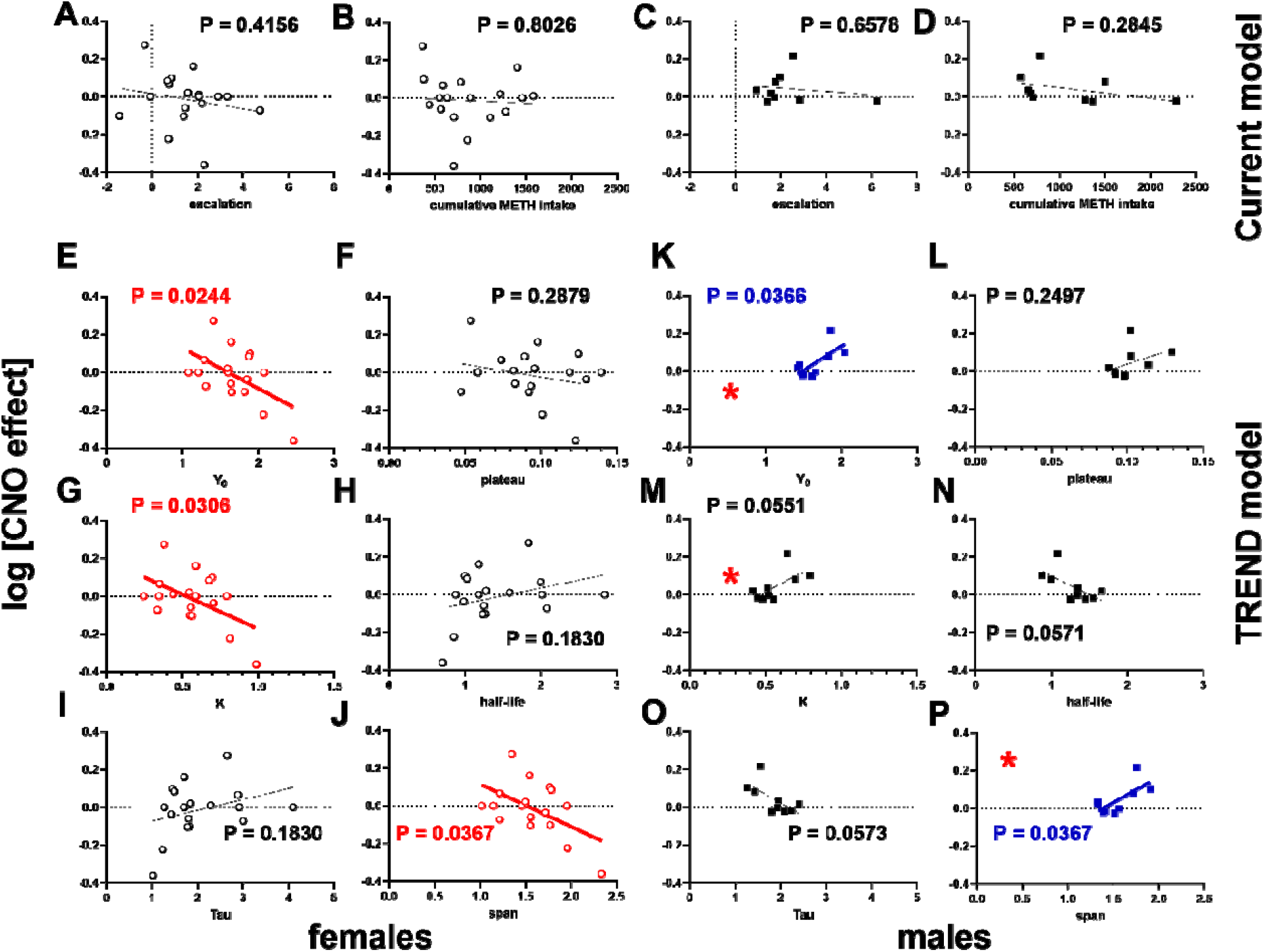
TREND model is more effective than the current model at detecting sex differences in the relationships between variables. We did not detect any sex differences in the gradient (slope) of the relationship between CNO effect (y-axis) and 1. escalation and 2. Cumulative METH intake for females (Fig A, B) and males (Fig C, D). However, there are interesting points to note about the variables derived from TREND model. For females, there are significant relationships between CNO effect (y-axis) and Y_0_ (Fig E), K (Fig G) and span (Fig J). A similar trend is observed in males, though the relationships are in the opposite direction (see Fig K, M and P). There are significant differences between males and females with regards to the highlighted relationships (see * insert in male graphs for difference between males and females). In summary there are significant sex differences in the relationships between variables derived from TREND, but not the current model, and CNO effect. P values are written into the graphs or * which represents P < 0.05.

There were no detected sex differences for any relationships between variables for the current model-derived variables. For comparisons between males and females for the relationship between escalation and CNO effect, we detected no sex differences (F 1, 23 = 0.1008, P = 0.7538). Similarly no sex differences for the relationship between cumulative METH intake and CNO effect (F 1, 23 = 0.08879, P = 0.7684). Interestingly, comparisons of the slopes of the relationship(s) between CNO effect and Y_0_, K and span revealed significant differences between males and females. For males versus females, the P values for the relationships between Y_0_ and CNO effect, K and CNO effect, span and CNO effect were P = 0.0221 (F 1, 23 = 6.026), P = 0.0300 (F 1, 23 = 5.352) and P = 0.0213 (F 1, 23 = 6.110), respectively. The comparisons, between males and females, of the slopes of the relationships between CNO effect and plateau, half-life and Tau returned P values of 0.2731 (F 1, 23 = 1.261), 0.1101 (F 1, 23 = 2.763), and 0.1102 (F 1, 23 = 2.761), respectively.

When we grouped our METH takers, males and females, into high versus low takers based on median split of the cumulative METH intake during the training phase (Fig 5A, current model) and assessed the effect of CNO on METH intake using 2-way ANOVA, we did not detect a SEX × group interaction (Fig 5C, F 1, 23 = 0.008313, P = 0.9281). However, when we grouped our subjects into high versus low behavioral outcome based on median split analysis of the variable ‘span’ (Fig 5B) derived from the TREND model (span is Y_0_ – plateau, see Fig 2C), we detected a SEX × group interaction (Fig 5D, F 1, 23 = 4.668, P = 0.0414). We explored groups of high and low escalation for males and females, but there was no SEX × group interaction with regards to escalation (F 1, 23 = 0.9500, P = 0.3398). We conducted the 2-way ANOVA and determined there was no SEX × group interaction for Y_0_ (F 1, 23 = 2.308, P = 0.1424) and K (F 1, 23 = 1.606, P = 0.2178).

**Figure 5:**
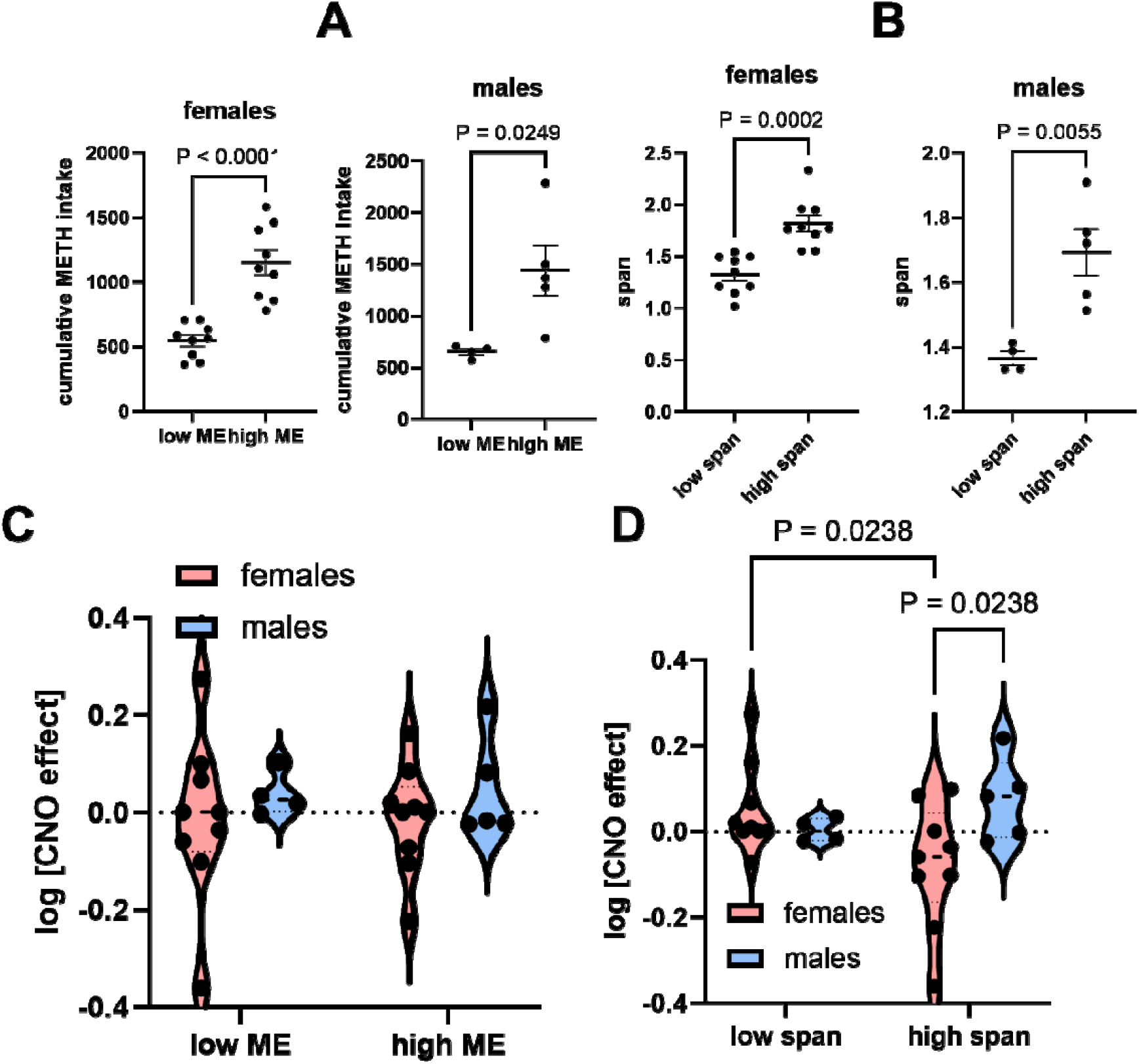
TREND model reveals METH experience-dependent sex differences; the current model did not detect any interactions. We conducted median split of cumulative METH intake (current model variable) for males and females to give rise to high and low METH experience groups (high ME, low ME, Fig A) and we did the same for span (Fig B). 2-way ANOVA with factors (SEX: males and females) and group (high and low responders determined by median split of cumulative METH intake in the current model) revealed no SEX × group interaction (P > 0.05, Fig C). Interestingly, median split of the TREND variable span to high and low span for males and females, followed by 2-way ANOVA of CNO effect (dependent variable) revealed a SEX × group interaction (Fig D, F 1, 23 = 4.668, P = 0.0414, Fig D) with uncorrected Fischer’s LSD showing that there were differences between males and females in the high span group (P = 0.0238) but not the low span group (p = 0.4905, Fig D). Interestingly also there were significant differences between the females in the low versus high span group (P = 0.0238, Fig D). Our analysis suggests that sex differences can be experience-dependent.

We wanted to determine if these new variables that correlated with CNO effects on METH and revealed sex differences were related to any of the current model variables. We determined if escalation and cumulative METH intake were related to Y_0_, plateau, K, half-life, Tau and span for males and females. We did not observe any relationships between current variables and TREND variables – these variables are dissimilar for males and females (P > 0.05, Fig 6). We determined that, for males and females, escalation was also unrelated to any of the TREND variables (data not shown).

**Figure 6:**
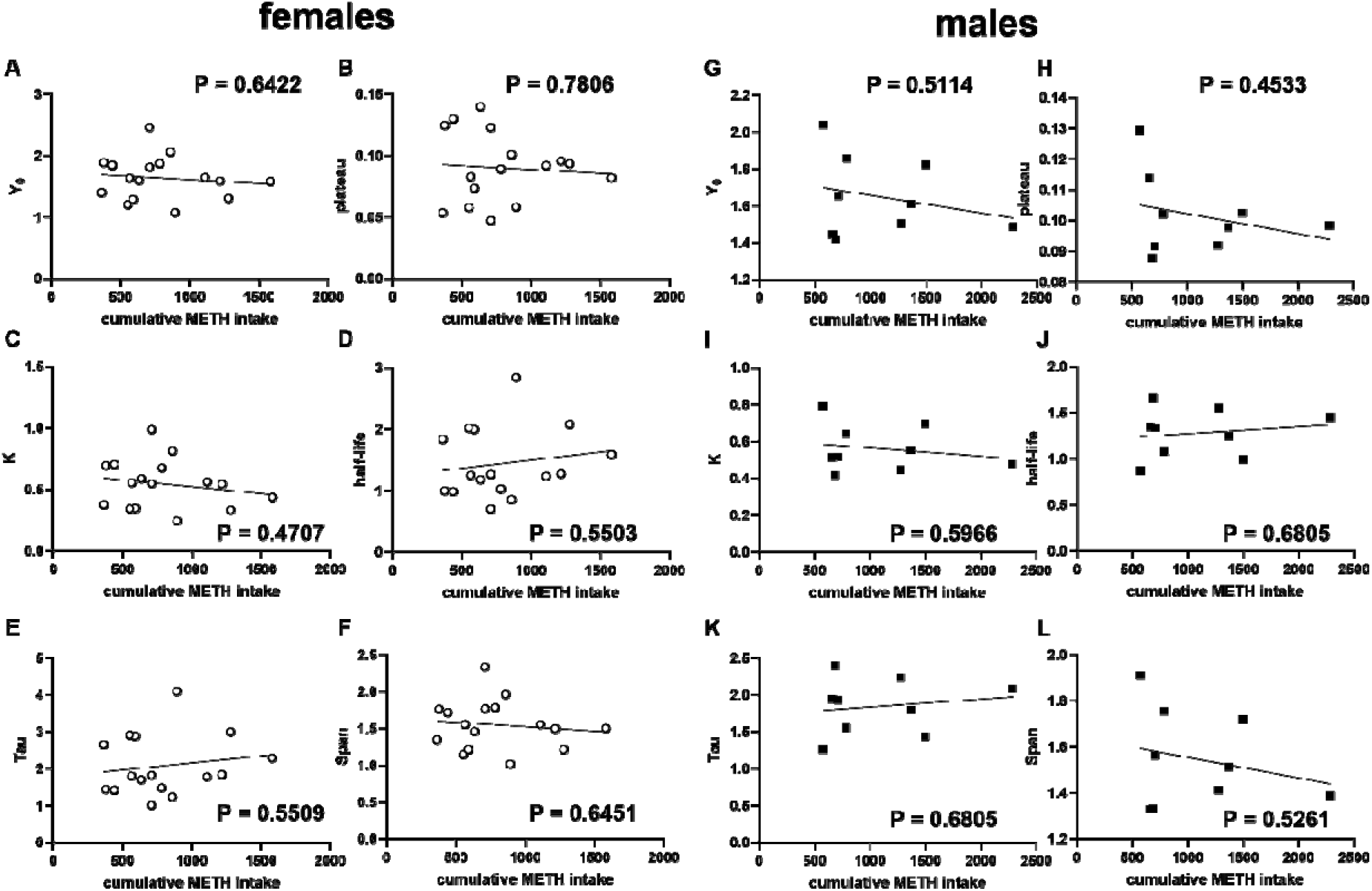
Variables from the current model and TREND do not overlap – there are no relationships between them. For females (Fig A-F) and males (Fig G-L), we examined relationships (correlations) between cumulative METH intake on the x-axis (current model) and all TREND model variables on the y-axis (see axis labels for appropriate variables). There were no significant relationships observed (P values are written into the graphs).

Overall, TREND model reveals sex differences in the effect of dorsal striatum dopamine D1 receptor activity inhibition on METH SA in a group of males and females that previously expressed high span but not low span. The current model was ineffective in detecting experience or sex differences in in the effect of dorsal striatum dopamine D1 receptor activity inhibition on METH SA, as reported previously (Job et al., 2020).

## Discussion

For this study, we wanted to address a gap in knowledge with regards to the role of dorsal striatal dopamine D1 receptor systems in METH self-administration and sex differences in this mechanism. Based on the literature, our hypothesis was that this role would be METH experience-dependent, with sex differences. We also hypothesized that one of the reasons for a lack of clarity in the above was because we were currently using current models for obtaining data that were not very sensitive. By conceptualizing ‘experience’ as a time-related continuum within which dynamic interactions between drug intake and cumulative drug intake occur, we developed a new exponential model termed the TREND model. We tested our hypothesis above using both the current model and the TREND model and we realized remarkably interesting results. We confirmed our hypothesis, but with the TREND model and not the current model.

TREND model-derived variables were more relevant to sex differences in the role of dorsal striatal dopamine D1 receptor systems in METH self-administration than the variables (escalation, cumulative METH intake) used in the current model. The TREND model determined that even through there was not an overall effect (average) of CNO on METH self-administration, CNO exerted significantly different effects on females with less span than females with more span (experience-dependent effects of dorsal striatal dopamine D1 receptor systems). TREND model also revealed that CNO exerted significantly different effects in males and females with high span (sex dependent effects of dorsal striatal dopamine D1 receptor systems). The current model used in the field was not able to determine any of the above. CNO effects were related to prior METH self-administration behavioral variables (Y_0_, K and span) derived from the TREND model. CNO effects were unrelated to any variables obtained from the current model. Importantly, TREND model revealed variables that were distinct from the ones currently used in the field. TREND is more effective than the current model in identifying sex and experience-dependent differences in the role of dorsal striatal dopamine D1 receptor systems in METH self-administration.

There are limitations to this study, see (Job et al., 2020). Firstly, DREADDs injected into the dorsal striatum were not localized to a specific compartment within this structure. The dorsomedial compartment of the dorsal striatum has been linked previously to initiation of drug taking activity, while the dorsolateral compartment of the dorsal striatum is thought to be more engaged in the transitioning to habitual drug taking behavior (Dalley and Everitt, 2009; Everitt and Robbins, 2013, 2005; Lüscher et al., 2020). As a result of non-specific non-subregion defined dorsal striatum dopamine D1 receptor-expressing neuron activity inhibition, we cannot delineate what specific function the dorsal striatum subregions may play in METH self-administration. More work needs to be done to clarify this. Another limitation has to do with CNO as a chemogenetic ligand. CNO is known to be converted to clozapine which can exert confounding pharmacological effects (Gomez et al., 2017; Ilg et al., 2018; MacLaren et al., 2016; Manvich et al., 2018; Neff et al., 2006; Padovan-Hernandez and Knackstedt, 2018; Rodd et al., 2022; Tanda et al., 2015; Ufer et al., 1999).

In summary, the dorsal striatal dopamine D1 receptor systems play a role in METH self-administration in a sex and experience-dependent manner. However, this was revealed by a new TREND model and not the current model. This report informs us, on the one hand, that we must account for prior drug experience (drug experience-time complex) when assessing the role of neurochemical systems in drug-related behavior and/or sex differences. But what is most important is that we have, for the first time, described an alternative model to the current model for more advanced studies, including with respect to understanding experience-related and sex-related differences. The implication may be that the currently used methods in the field may represent a less sensitive model for detecting experience-related and sex-related differences in drug use but also in other behavioral studies. Thus, our work may advance the field.

## Acknowledgements

The authors wish to acknowledge Dr. Jean Lud Cadet in whose laboratory MOJ conducted the behavioral experiments. The authors also acknowledge Dr. Atul Daiwile and Mr. Michael Chojnacki who contributed to the behavioral experiments. All authors contributed to data analysis and to the writing of the manuscript. MOJ designed and conducted the behavioral experiments and statistical analysis. This work was funded by the Department of Health and Human Services/National Institutes of Health/National Institute on Drug Abuse/Intramural Research Program, Baltimore, MD, USA [grant #-DA000552]. This work was also supported by the Francis Lax Fund for Faculty Development at Rowan University. This work was also supported by startup funds from Rowan University, Camden, New Jersey.

## Disclosures

InduMithra Madhuranthakam has no conflicts of interest to declare. Dr. Martin O Job has no conflicts of interest to declare.

## Notes

### Competing Interest Statement

The authors have declared no competing interest.

## References

Avchalumov, Y., Trenet, W., Piña-Crespo, J., Mandyam, C., 2020. Sch23390 reduces methamphetamine self-administration and prevents methamphetamine-induced striatal ltd. Int. J. Mol. Sci. 21, 1–16. 10.3390/ijms21186491

Bachtell, R.K., Larson, T.A., Winkler, M.C., 2023. Adenosine receptor stimulation inhibits methamphetamine-associated cue seeking. J. Psychopharmacol. 37, 192–203. 10.1177/02698811221147157

Belin, D., Everitt, B.J., 2008. Cocaine seeking habits depend upon dopamine-dependent serial connectivity linking the ventral with the dorsal striatum. Neuron 57, 432–441. 10.1016/J.NEURON.2007.12.019

Chojnacki, M.R., Jayanthi, S., Cadet, J.L., 2020. Methamphetamine pre-exposure induces steeper escalation of methamphetamine self-administration with consequent alterations in hippocampal glutamate AMPA receptor mRNAs 889.

Culbertson, C., De La Garza, R., Costello, M., Newton, T.F., 2009. Unrestricted access to methamphetamine or cocaine in the past is associated with increased current use. Int. J. Neuropsychopharmacol. 12, 677–685. 10.1017/S1461145708009668

Daiwile, A.P., Jayanthi, S., Cadet, J.L., 2022. Sex differences in methamphetamine use disorder perused from pre-clinical and clinical studies: Potential therapeutic impacts 137, 104674.

Daiwile, A.P., Jayanthi, S., Cadet, J.L., 2021. Sex- and Brain Region-specific Changes in Gene Expression in Male and Female Rats as Consequences of Methamphetamine Self-administration and Abstinence. Neuroscience 452, 265–279.

Dalley, J.W., Everitt, B.J., 2009. Dopamine receptors in the learning, memory and drug reward circuitry. Semin. Cell Dev. Biol. 20, 403–410. 10.1016/J.SEMCDB.2009.01.002

Dubois, A., Savasta, M., Curet, O., Scatton, B., 1986. Autoradiographic distribution of the D1 agonist [3H]SKF 38393, in the rat brain and spinal cord. Comparison with the distribution of D2 dopamine receptors. Neuroscience 19, 125–137. 10.1016/0306-4522(86)90010-2

Everitt, B.J., Robbins, T.W., 2013. From the ventral to the dorsal striatum: devolving views of their roles in drug addiction. Neurosci. Biobehav. Rev. 37, 1946–54. 10.1016/j.neubiorev.2013.02.010

Everitt, B.J., Robbins, T.W., 2005. Neural systems of reinforcement for drug addiction: From actions to habits to compulsion. Nat. Neurosci. 10.1038/nn1579

Gomez, J.L., Bonaventura, J., Lesniak, W., Mathews, W.B., Sysa-Shah, P., Rodriguez, L.A., Ellis, R.J., Richie, C.T., Harvey, B.K., Dannals, R.F., Pomper, M.G., Bonci, A., Michaelides, M., 2017. Chemogenetics revealed: DREADD occupancy and activation via converted clozapine. Science (80-.). 357, 503–507. 10.1126/science.aan2475

Holtz, N.A., Lozama, A., Prisinzano, T.E., Carroll, M.E., 2012. Reinstatement of methamphetamine seeking in male and female rats treated with modafinil and allopregnanolone. Drug Alcohol Depend. 120, 233–237. 10.1016/J.DRUGALCDEP.2011.07.010

Ilg, A.K., Enkel, T., Bartsch, D., Bähner, F., 2018. Behavioral Effects of Acute Systemic Low-Dose Clozapine in Wild-Type Rats: Implications for the Use of DREADDs in Behavioral Neuroscience. Front. Behav. Neurosci. 12. 10.3389/FNBEH.2018.00173

Job, M.O., Chojnacki, M.R., Daiwile, A.P., Cadet, J.L., 2020. Chemogenetic Inhibition of Dopamine D1-expressing Neurons in the Dorsal Striatum does not alter Methamphetamine Intake in either Male or Female Long Evans Rats. Neurosci. Lett. 729. 10.1016/j.neulet.2020.134987

Johansen, A., McFadden, L.M., 2017. The neurochemical consequences of methamphetamine self-administration in male and female rats. Drug Alcohol Depend. 178, 70–74. 10.1016/J.DRUGALCDEP.2017.04.011

Kazahaya, Y., Akimoto, K., Otsuki, S., 1989. Subchronic methamphetamine treatment enhances methamphetamine-or cocaine-induced dopamine efflux in vivo. Biol. Psychiatry 25, 903–912. 10.1016/0006-3223(89)90270-9

Kearns, A.M., Siemsen, B.M., Hopkins, J.L., Weber, R.A., Scofield, M.D., Peters, J., Reichel, C.M., 2022. Chemogenetic inhibition of corticostriatal circuits reduces cued reinstatement of methamphetamine seeking. Addict. Biol. 27. 10.1111/ADB.13097

Kreisler, A.D., Terranova, M.J., Somkuwar, S.S., Purohit, D.C., Wang, S., Head, B.P., Mandyam, C.D., 2020. In vivo reduction of striatal D1R by RNA interference alters expression of D1R signaling-related proteins and enhances methamphetamine addiction in male rats. Brain Struct. Funct. 225, 1073–1088. 10.1007/s00429-020-02059-w

Lewandowski, S.I., Hodebourg, R., Wood, S.K., Carter, J.S., Nelson, K.H., Kalivas, P.W., Reichel, C.M., 2023. Matrix metalloproteinase activity during methamphetamine cued relapse. Addict. Biol. 28. 10.1111/ADB.13279

Lin, H., Olaniran, A., Garmchi, S., Firlie, J., Rincon, N., Li, X., 2024. The estrous cycle has no effect on incubation of methamphetamine craving and associated Fos expression in dorsomedial striatum and anterior intralaminar nucleus of thalamus. Addict. Neurosci. 11. 10.1016/J.ADDICN.2024.100158

Lorrain, D.S., Arnold, G.M., Vezina, P., 2000. Previous exposure to amphetamine increases incentive to obtain the drug: Long-lasting effects revealed by the progressive ratio schedule. Behav. Brain Res. 107, 9–19. 10.1016/S0166-4328(99)00109-6

Lüscher, C., Robbins, T., Everitt, B., 2020. The transition to compulsion in addiction. Nat. Rev. Neurosci. 21, 247–263. 10.1038/S41583-020-0289-Z

MacLaren, D.A.A., Browne, R.W., Shaw, J.K., Radhakrishnan, S.K., Khare, P., España, R.A., Clark, S.D., 2016. Clozapine N-oxide administration produces behavioral effects in long-evans rats: Implications for designing DREADD experiments. eNeuro 3. 10.1523/ENEURO.0219-16.2016

Manvich, D.F., Webster, K.A., Foster, S.L., Farrell, M.S., Ritchie, J.C., Porter, J.H., Weinshenker, D., 2018. The DREADD agonist clozapine N-oxide (CNO) is reverse-metabolized to clozapine and produces clozapine-like interoceptive stimulus effects in rats and mice. Sci. Rep. 8. 10.1038/S41598-018-22116-Z

McFadden, L.M., Hadlock, G.C., Allen, S.C., Vieira-Brock, P.L., Stout, K.A., Ellis, J.D., Hoonakker, A.J., Andrenyak, D.M., Nielsen, S.M., Wilkins, D.G., Hanson, G.R., Fleckenstein, A.E., 2012. Methamphetamine self-administration causes persistent striatal dopaminergic alterations and mitigates the deficits caused by a subsequent methamphetamine exposure. J. Pharmacol. Exp. Ther. 340, 295–303. 10.1124/jpet.111.188433

Mcfadden, L.M., Hanson, G.R., Fleckenstein, A.E., 2013. The effects of methamphetamine self-administration on cortical monoaminergic deficits induced by subsequent high-dose methamphetamine administrations. Synapse 67, 875–881. 10.1002/SYN.21696

McFadden, L.M., Vieira-Brock, P.L., Hanson, G.R., Fleckenstein, A.E., 2015. Prior methamphetamine self-administration attenuates the dopaminergic deficits caused by a subsequent methamphetamine exposure. Neuropharmacology 93, 146–154.

Neff, N.H., Wemlinger, T.A., Duchemin, A.M., Hadjiconstantinou, M., 2006. Clozapine modulates aromatic L-amino acid decarboxylase activity in mouse striatum. J. Pharmacol. Exp. Ther. 317, 480–487. 10.1124/JPET.105.097972

Okita, K., Morales, A.M., Dean, A.C., Johnson, M.C., Lu, V., Farahi, J., Mandelkern, M.A., London, E.D., 2018. Striatal dopamine D1-type receptor availability: No difference from control but association with cortical thickness in methamphetamine users. Mol. Psychiatry 23, 1320–1327. 10.1038/MP.2017.172

Oliver, R.J., Purohit, D.C., Kharidia, K.M., Mandyam, C.D., 2019. Transient Chemogenetic Inhibition of D1-MSNs in the Dorsal Striatum Enhances Methamphetamine Self-Administration. Brain Sci. 9. 10.3390/brainsci9110330

Orio, L., Wee, S., Newman, A.H., Pulvirenti, L., Koob, G.F., 2010. The dopamine D3 receptor partial agonist CJB090 and antagonist PG01037 decrease progressive ratio responding for methamphetamine in rats with extended-access. Addict. Biol. 15, 312–323. 10.1111/J.1369-1600.2010.00211.X

Padovan-Hernandez, Y., Knackstedt, L.A., 2018. Dose-dependent reduction in cocaine-induced locomotion by Clozapine-N-Oxide in rats with a history of cocaine self-administration. Neurosci. Lett. 674, 132–135. 10.1016/J.NEULET.2018.03.045

Pena-Bravo, J.I., Penrod, R., Reichel, C.M., Lavin, A., 2019. Methamphetamine self-administration elicits sex-related changes in postsynaptic glutamate transmission in the prefrontal cortex. eNeuro 6. 10.1523/ENEURO.0401-18.2018

Pierre, P.J., Vezina, P., 1998. D1 dopamine receptor blockade prevents the facilitation of amphetamine self-administration induced by prior exposure to the drug. Psychopharmacology (Berl). 138, 159–166. 10.1007/s002130050658

Pittenger, S.T., Chou, S., Murawski, N.J., Barrett, S.T., Loh, O., Duque, J.F., Li, M., Bevins, R.A., 2021. Female rats display higher methamphetamine-primed reinstatement and c-Fos immunoreactivity than male rats. Pharmacol. Biochem. Behav. 201. 10.1016/J.PBB.2020.173089

Rodd, Z.A., Engleman, E.A., Truitt, W.A., Burke, A.R., Molosh, A.I., Bell, R.L., Hauser, S.R., 2022. CNO Administration Increases Dopamine and Glutamate in the Medial Prefrontal Cortex of Wistar Rats: Further Concerns for the Validity of the CNO-activated DREADD Procedure. Neuroscience 491, 176–184. 10.1016/J.NEUROSCIENCE.2022.03.028

Savasta, M., Dubois, A., Scatton, B., 1986. Autoradiographic localization of D1 dopamine receptors in the rat brain with [3H]SCH 23390. Brain Res. 375, 291–301. 10.1016/0006-8993(86)90749-3

Segal, D.S., Kuczenski, R., O’Neil, M.L., Melega, W.P., Cho, A.K., 2003. Escalating dose methamphetamine pretreatment alters the behavioral and neurochemical profiles associated with exposure to a high-dose methamphetamine binge. Neuropsychopharmacology 28, 1730–1740. 10.1038/sj.npp.1300247

Shishido, T., Watanabe, Y., Suzuki, H., Kato, K., Niwa, S., Hanoune, J., Matsuoka, I., 1997. Effects of repeated methamphetamine administration on dopamine D1 receptor, D2 receptor and adenylate cyclase type V mRNA levels in the rat striatum. Neurosci. Lett. 222, 175–178. 10.1016/S0304-3940(97)13376-6

Somkuwar, S.S., Fannon, M.J., Head, B.P., Mandyam, C.D., 2016. Methamphetamine reduces expression of caveolin-1 in the dorsal striatum: Implication for dysregulation of neuronal function. Neuroscience 328, 147–56. 10.1016/j.neuroscience.2016.04.039

Tanda, G., Valentini, V., De Luca, M.A., Perra, V., Serra, G. Pietro, Di Chiara, G., 2015. A systematic microdialysis study of dopamine transmission in the accumbens shell/core and prefrontal cortex after acute antipsychotics. Psychopharmacology (Berl). 232, 1427–1440. 10.1007/S00213-014-3780-2

Tong, J., Ross, B., Schmunk, G., Peretti, F., Kalasinsky, K., Furukawa, Y., Ang, L., Aiken, S., Wickham, D., Kish, S., 2003. Decreased striatal dopamine D1 receptor-stimulated adenylyl cyclase activity in human methamphetamine users. Am. J. Psychiatry 160, 896–903. 10.1176/APPI.AJP.160.5.896

Ufer, M., Dadmarz, M., Vogel, W.H., 1999. Voluntary consumption of amphetamine, cocaine, ethanol and morphine by rats as influenced by a preceding period of forced drug intake and clozapine. Pharmacology 58, 285–291. 10.1159/000028293

Vanderschuren, L.J.M.J., Everitt, B.J., 2004. Drug seeking becomes compulsive after prolonged cocaine self-administration. Science 305, 1017–1019. 10.1126/SCIENCE.1098975

Veeneman, M.M.J., Broekhoven, M.H., Damsteegt, R., Vanderschuren, L.J.M.J., 2012. Distinct contributions of dopamine in the dorsolateral striatum and nucleus accumbens shell to the reinforcing properties of cocaine. Neuropsychopharmacology 37, 487–498. 10.1038/npp.2011.209

Venniro, M., Zhang, M., Shaham, Y., Caprioli, D., 2017. Incubation of Methamphetamine but not Heroin Craving after Voluntary Abstinence in Male and Female Rats. Neuropsychopharmacology 42, 1126–1135. 10.1038/npp.2016.287

Vezina, P., Pierre, P.J., Lorrain, D.S., 1999. The effect of previous exposure to amphetamine on drug-induced locomotion and self-administration of a low dose of the drug. Psychopharmacology (Berl). 147, 125–134. 10.1007/s002130051152

Wang, J.Q., McGinty, J.F., 1995. Differential Effects of D1 and D2 Dopamine Receptor Antagonists on Acute Amphetamine or Methamphetamine Induced Up Regulation of zif/268 mRNA Expression in Rat Forebrain. J. Neurochem. 65, 2706–2715. 10.1046/j.1471-4159.1995.65062706.x

Wee, S., Wang, Z., Woolverton, W.L., Pulvirenti, L., Koob, G.F., 2007. Effect of aripiprazole, a partial dopamine D2 receptor agonist, on increased rate of methamphetamine self-administration in rats with prolonged session duration. Neuropsychopharmacology 32, 2238–2247. 10.1038/SJ.NPP.1301353

Westbrook, S.R., Dwyer, M.R., Cortes, L.R., Gulley, J.M., 2020. Extended access self-administration of methamphetamine is associated with age- and sex-dependent differences in drug taking behavior and recognition memory in rats. Behav. Brain Res. 390. 10.1016/J.BBR.2020.112659

Yoshida, H., Ohno, M., Watanabe, S., 1995. Roles of dopamine D1 receptors in striatal fos protein induction associated with methamphetamine behavioral sensitization in rats. Brain Res. Bull. 38, 393–397. 10.1016/0361-9230(95)02005-C

